# Mapping research on Indigenous peoples, traditional knowledge, and biodiversity conservation in the Amazon: gaps and Indigenous knowledge co-production

**DOI:** 10.64898/2026.04.13.718171

**Authors:** Jady Vivia Almeida da Silva Santos, Francieli F. Bomfim, Josinete Sampaio Monteneles, Mayerly Alexandra Guerrero-Moreno, Yan Campioni, Everton Cruz da Silva, Joás Silva Brito, José Max Barbosa Oliveira-Junior, Kiarasy Kaiabi Panara, Korakoko Panara, Sewa Panara, Sakre Panara, Karapow Panara, Kwakore Panara, Sopoa Panara, Nhasykiati Panara, Pâssua Pri Panara, Pente Panara, Tepakriti Panara, Tyson Ferreira-Sateré, Amanda Kumaruara, Yuri Kuikuro, Antonio Ramyllys Oliveira Costa, Lais Sarlo, Bruno Coutinho, Renata Pinheiro, Rafael de Araújo, Marcos Pérsio Dantas Santos, Ana Cristina Mendes-Oliveira, Gleomar Fabiano Maschio, Erival Gonçalves Prata, Bruno Marangoni Martinelli, Isabella Millena Alves Evangelista, Domingos de Jesus Rodrigues, Karina Dias-Silva, Luciano Fogaça de Assis Montag, Thaisa Sala Michelan, Leandro Juen

## Abstract

1. Indigenous peoples play a central role in biodiversity knowledge and conservation, yet their participation in scientific research remains underrepresented. Understanding how Indigenous peoples, traditional knowledge, and Indigenous territories are portrayed in the scientific literature is essential for developing more equitable and culturally grounded conservation strategies.
2. We conducted a bibliometric analysis of 94 articles on biodiversity conservation in the Amazon, published between 1997 and 2025, indexed in Scopus and Web of Science. We examined temporal trends, geographic distribution, institutional leadership, Indigenous co-authorship, focal ecosystems and taxa, and the main contributions attributed to Indigenous peoples. Indigenous perspectives were integrated into this analysis through a participatory approach.
3. Scientific production increased after 2010. Research leadership remains concentrated in institutions from the Global North, even though Brazil, Ecuador, and Peru were the most frequently studied countries. Indigenous co-authorship was identified in only 6.4 % of the studies. Most studies focused on plants, mammals, and birds, whereas aquatic environments and groups such as insects, amphibians, and reptiles received comparatively less attention. The main contributions attributed to Indigenous peoples were related to community-based monitoring and management (41.48%) and cultural practices and traditional ecological knowledge (38.19%).
4. These findings show that Indigenous peoples are widely recognized as knowledge holders and conservation actors, but are still rarely included as authors or research partners. Our study highlights persistent geographic, epistemic, and collaborative asymmetries in Amazonian biodiversity research. Conservation science and policy will be stronger, fairer, and more effective when they move beyond documenting Indigenous knowledge towards supporting Indigenous leadership, equitable partnerships, and inclusive co-production of knowledge.

## 1. Introduction

Biodiversity loss is one of the most critical environmental challenges of our time, accelerating species decline, reducing essential ecosystem services, and undermining human well-being (Hogue & Breon, 2022; Díaz et al., 2019; Ceballos et al., 2010). In megadiverse regions such as the Amazon, these threats are intensified by deforestation, agricultural expansion, and mining, which accelerate habitat degradation and loss while weakening communities that depend directly on natural resources for their livelihoods and sociocultural reproduction (Fearnside, 2017; Silva Junior et al., 2021; Joly et al., 2020; Da Silva et al., 2024; Brandão et al., 2022).

In this context, Indigenous peoples play a crucial role in conserving natural resources. Their ways of life, grounded in the multiple use of nature, are closely linked to ecological integrity and supported by knowledge systems deeply rooted in territory and biodiversity management, transmitted across generations, and reflecting integrated understandings of environmental dynamics and society-land relationships (Schmink et al., 2019; Levis et al., 2024; Da Silva et al., 2024; Berkes et al., 2000; Gadgil et al., 1993). These knowledge systems provide a foundation for participatory biodiversity management and conservation strategies in local communities and can also inform broader conservation models (Leret-Câmara et al., 2019; Batista et al., 2020; Varese et al., 2021).

Cultural practices, traditional farming systems, and ecological perceptions help maintain essential ecological processes, support ecosystem services, enhance socioecological resilience, and sustain evolutionary processes (Levis et al., 2024; Folhes et al., 2025). A substantial share of global biodiversity occurs in lands traditionally managed by Indigenous peoples, and preserved Amazonian habitats, including primary forests, have been shaped and managed by these populations over centuries (Garnett et al., 2018; Barlow et al., 2012; Nascimento et al., 2024).

Despite this recognition, reinforced by frameworks such as Brazil’s National Policy for Territorial and Environmental Management of Indigenous Lands (PNGATI), a persistent gap remains between institutional discourse and conservation practice (Guimarães et al., 2014; Gannon et al., 2019; Witter & Satterfield, 2019; Dawson et al., 2023). Although these initiatives emphasize incorporating traditional knowledge and the active participation of Indigenous communities in conservation policy, this recognition has not yet translated into effective inclusion in scientific research and decision-making (Corrigan et al., 2018; Osborne et al., 2024; Berkes et al., 2000). The challenge of intercultural dialogue between Indigenous sciences and academic science is reflected in the scarcity of studies on ecological knowledge in Indigenous territories and in the limited participation of Indigenous peoples in research processes (Carvalho et al., 2023). As a result, approaches that privilege Western science alone often overlook the equitable inclusion of traditional knowledge and produce conservation policies that are poorly aligned with local realities (Levis et al., 2024; Da Silva et al., 2024).

Although recent initiatives have demonstrated the success of community-based ecological monitoring and Indigenous leadership in conservation, relationships between knowledge systems remain shaped by power asymmetries (Constantino et al., 2008; Freitas et al., 2019; Moraes et al., 2024; Juruna et al., 2025; Sheil et al., 2015). These examples show that Indigenous participation is essential for Amazon conservation, as ecological perceptions, sustainable practices, and deep connections with forests are fundamental to maintaining biodiversity and ecological functioning (Levis et al., 2024; Santos et al., 2024).

Although dialogue between traditional and scientific knowledge is frequently invoked in international agreements, these encounters remain shaped by historically entrenched power relations rooted in epistemic inequalities and the hierarchical positioning of Western science (Martinelli et al., 2022; Lunetta et al., 2025). Concrete actions that value Indigenous leadership and respect Indigenous sciences are therefore essential, alongside public policies that safeguard Indigenous rights and strengthen conservation through collaboration between knowledge systems (Berkes, 2017; Díaz et al., 2019).

Against this backdrop, this study uses a bibliometric approach to examine how the scientific literature has linked biodiversity conservation to the traditional knowledge of Indigenous peoples in the Amazon and how Indigenous peoples are represented in the production of scientific knowledge on biodiversity conservation. We identify patterns, temporal trends, and gaps in the literature to map the thematic development of the field, recognize emerging directions, and inform evidence-based research agendas and public policies (Guerrero-Moreno et al., 2024; Da Silva et al., 2024; Oliveira-Junior et al., 2022).

Specifically, we ask: (i) How has scientific production on this topic changed over time? (ii) Which geographic areas and Indigenous territories have been most frequently studied? (iii) Which countries and institutions lead that research? (iv) How are institutional affiliations and Indigenous co-authorship represented in studies on biodiversity conservation in the Amazon? (v) What kinds of contributions are attributed to Indigenous communities? Furthermore, (vi) which ecosystems, taxa, and methods have been prioritized? By answering these questions, we aim to identify advances, inequalities, and opportunities to strengthen Indigenous leadership in global biodiversity conservation agendas.

## 2. Materials and Methods

### 2.1 Study scope and database construction

This study focused on the Pan-Amazon region, which encompasses the world’s largest tropical forest and one of its richest centers of biodiversity (Góes et al., 2025). This geographical region is an important reservoir of ancestral knowledge held by Indigenous peoples whose ways of life are closely linked to the conservation of local ecosystems. We used a bibliometric approach to analyze the international scientific literature, allowing the identification of publication trends, authorship profiles, geographic distribution and contributions attributed to Indigenous peoples over time. Literature searches were conducted in the Scopus and Web of Science, selected for their broad multidisciplinary coverage and representation of global scientific production (Da Silva et al., 2024).

In both databases, we applied the search string (Indigenous AND Biodiversity AND Conservation AND Amazon*) to title, abstract, and keywords. The truncation operator (*) was used to capture variations such as “Amazonia” and “Amazonian”, and the Boolean operator AND ensured the inclusion of all thematic components.

### 2.2 Inclusion and exclusion criteria

To ensure consistency, we included only original research articles published in peer-reviewed journals. We excluded reviews, book chapters, conference abstracts, notes, and theses. The temporal scope ranged from 1997 to 2025, corresponding to the earliest record identified. Searches were conducted on 28 August 2025. No language restrictions were applied to include diverse regional and cultural perspectives (Walpole et al., 2019; Guerrero-Moreno et al., 2024) (Figure 1).

**Figure 1.**
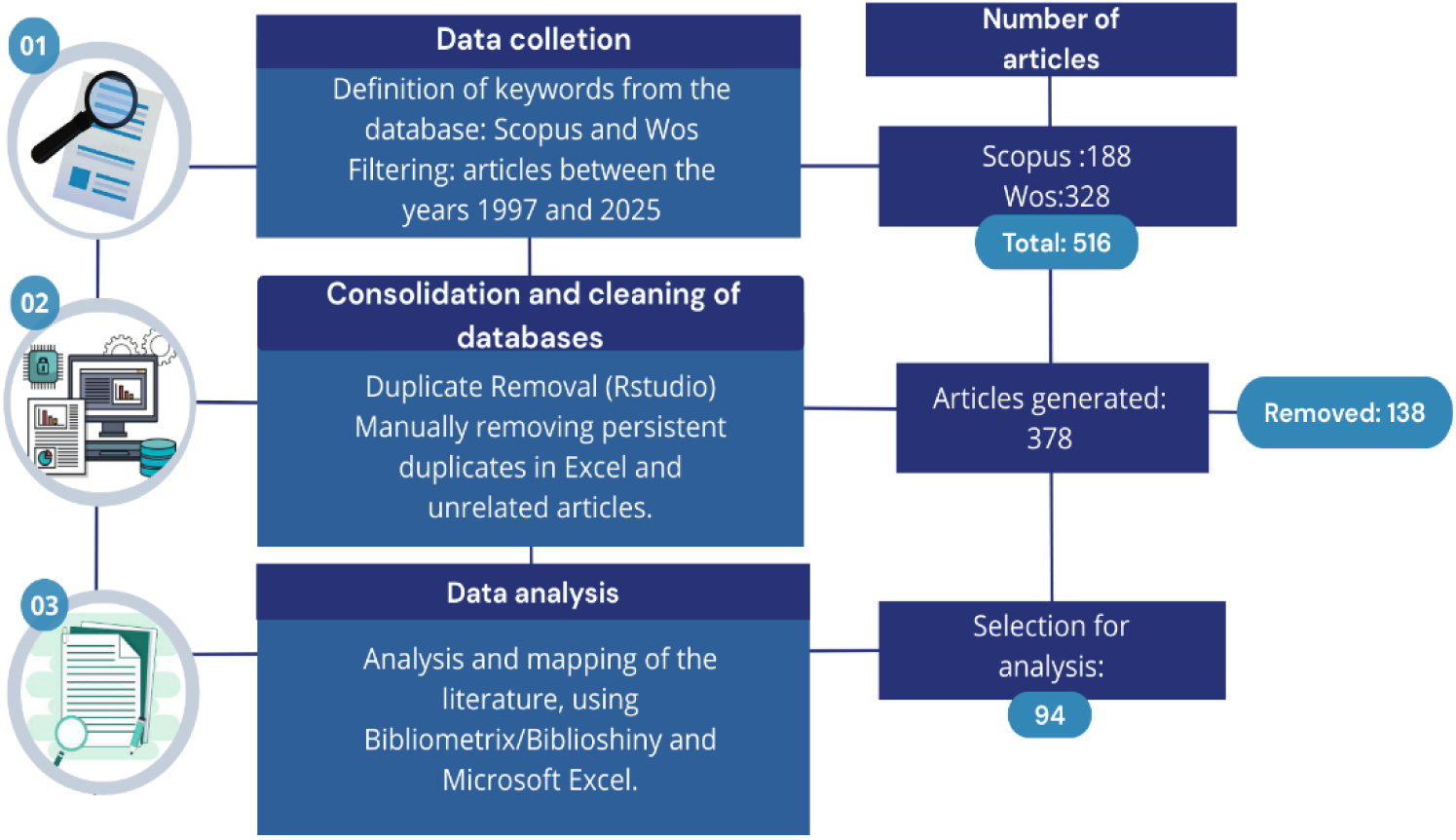
General flowchart of the methodology used for the search, consolidation, cleaning, and analysis of articles.

### 2.3 Data organization and categorization

We screened titles, abstracts, keywords, and methods to assess the study objectives (Costa et al., 2025). Based on this screening, we extracted structured information across five analytical axes:

1. Indigenous participation: Whether Indigenous peoples were involved in conservation-related actions and strategies.
2. Types of contribution: classified into four main categories, adapted from Da Silva et al. (2024): (i) cultural practices and traditional ecological knowledge; (ii) management and sustainable resource use; (iii) community-based monitoring and co-production of knowledge; and (iv) habitat conservation and restoration.
3. Study area: classification of Indigenous Lands and other protected areas (e.g., National Parks, RESEX) (SNUC, 2000).
4. Indigenous co-authorship: presence and position of Indigenous authors in publications.
5. Taxa investigated: biological groups addressed in each study.

### 2.4 Integration of Indigenous perspectives and participatory approach

In addition to the bibliometric analysis, we incorporated a qualitative stage involving Indigenous researchers connected to the study contexts. This approach aimed to broaden interpretation through lived experience and support knowledge co-production. Preliminary results were shared with Indigenous collaborators from different Amazonian territories (Panará, Sateré-Mawé, Kumaruara, and Kuikuro) who reflected on: (i) Indigenous co-authorship; (ii) Global North leadership; (iii) research priorities; (iv) necessary changes in scientific practice.

Contributions were obtained through direct communication, including written feedback, videos, and dialogical exchanges. These perspectives were systematized descriptively and incorporated into the interpretation of results, preserving authorship and narrative integrity. This stage allowed for contextualising bibliometric patterns through underrepresented perspectives, strengthening an inclusive and reflexive research approach (Levis et al., 2024; Berkes et al., 2000).

Authorization to access Indigenous territories was granted by the Fundação Nacional dos Povos Indígenas (FUNAI) under administrative process No. 08620.012263/2023-71. The capture, collection, and transport of biological material were authorized by the Instituto Brasileiro do Meio Ambiente e dos Recursos Naturais Renováveis (IBAMA; SISBIO permit no. 4681-1) and approved by the Animal Use Ethics Committee of the Federal University of Pará (CEUA No. 8293020418).

### 2.5 Data analysis

Data were analyzed using descriptive statistics and graphical visualization. All analyses were conducted in RStudio (version 4.4.0) using the Bibliometrix/Biblioshiny packages (Aria & Cuccurullo, 2017). Figures were finalized using Canva.

## 3. Results and Discussion

### 3.1 Temporal patterns and trends in scientific production

Between 1997 and the early 2000s, only a small number of studies were identified, with sporadic records and no publications in some years. From 2007 onwards, scientific production began to increase modestly, with a first more pronounced rise in 2008 (Figure 2). Between 2012 and 2016, publication output remained relatively stable, ranging from two to four studies per year. These patterns suggest that although recognition of the importance of Indigenous peoples to biodiversity conservation has advanced, its incorporation into scientific literature has been gradual rather than linear.

**Figure 2.**
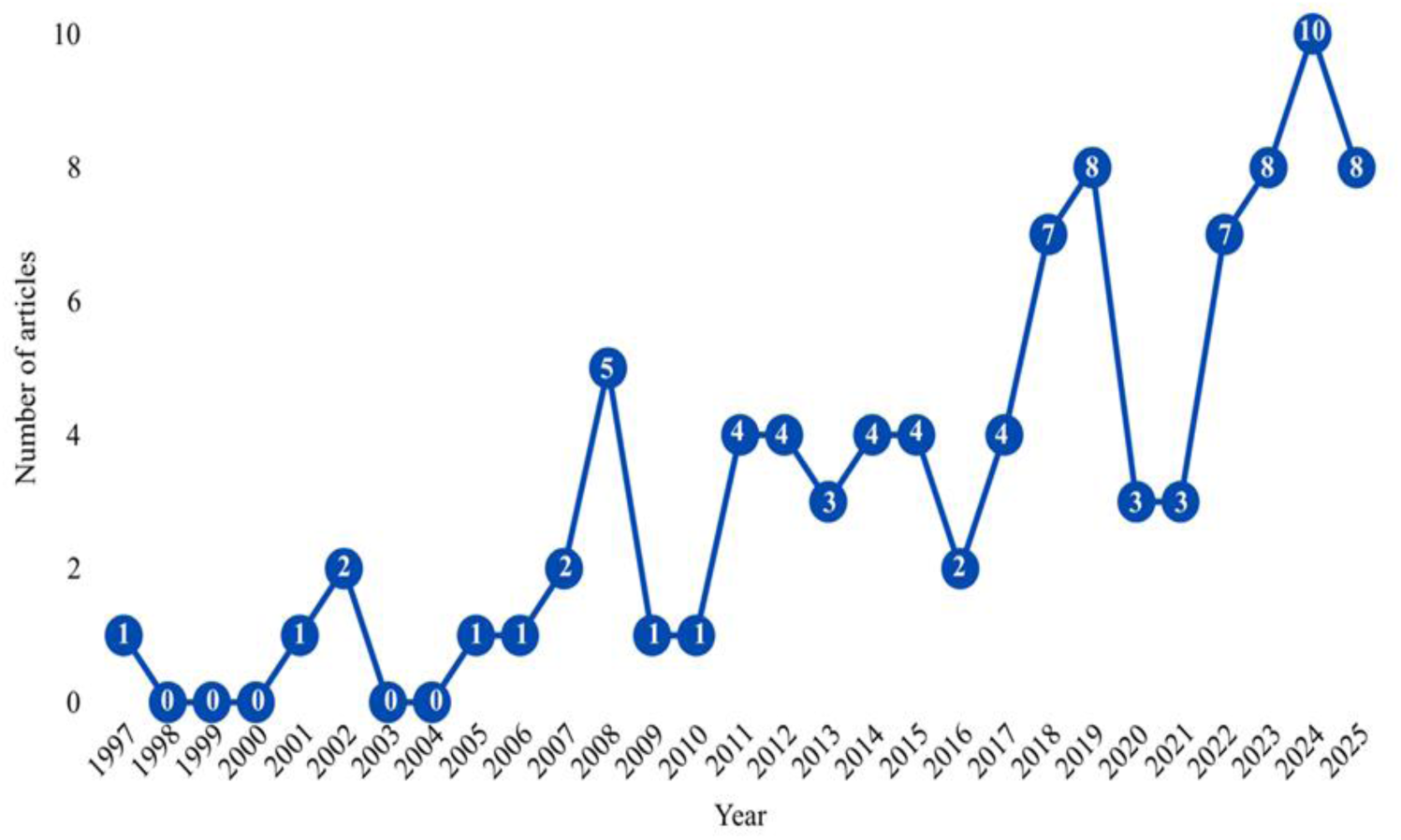
Annual scientific production of studies addressing the participation of Indigenous peoples in Amazon biodiversity conservation.

This temporal pattern indicates that growth in scientific production on the role of Indigenous communities in Amazon biodiversity conservation has coincided with political, institutional, and environmental milestones that increased the visibility of Indigenous leadership in conservation agendas. One important landmark was the 1992 United Nations Conference on Environment and Development (Earth Summit), whose Agenda 21 included a chapter dedicated to the role of Indigenous communities in environmental management and the sustainable use of natural resources (UN, 1992). This was followed by the strengthening of international agreements such as the Convention on Biological Diversity, which recognized traditional knowledge as central to biodiversity conservation and sustainable use (Dutfield, 2000).

The earliest studies identified in the 1990s reflect the growing recognition of Indigenous knowledge, particularly in interdisciplinary fields such as ethnobotany, ethnoecology, and ethnoichthyology (Athayde et al., 2025). Pioneering studies, such as Posey (1997), highlighted the relevance of traditional knowledge by demonstrating that Kayapó knowledge of plants was consistent with the principles of the Convention on Biological Diversity. A few years later, *Conservation and development alliances with the Kayapó of south-eastern Amazonia* (Zimmerman et al., 2001) assessed the ecological success achieved by the Kayapó in partnership with Conservation International (CI). The authors demonstrated the value of participatory monitoring strategies and of cooperation between Indigenous communities and conservation organisations, highlighting the active roles of these populations in environmental governance. Together, these early studies helped establish the importance of traditional knowledge in environmental science and laid methodological foundations for participatory approaches that later became important references for conservation research in Indigenous territories.

The increase in scientific production observed from 2018 onwards may be associated with the consolidation of international initiatives such as the Nagoya Protocol (2010), together with national policies aimed at valuing Indigenous knowledge, which created a more favorable environment for research foregrounding Indigenous peoples. The publication peak recorded in 2024 may reflect the recent strengthening of international debate on Indigenous roles in conservation, driven by events such as COP16 in Cali, Colombia, which emphasized the central role of Indigenous peoples in biodiversity conservation. It may also be linked to the launch of the Kuntari Katu Program, which seeks to expand Indigenous participation in climate negotiations and environmental public policy (MPI, 2023). Taken together, these milestones suggest that international and national agendas have played an important role in shaping the visibility and growth of scientific production on Indigenous peoples.

By contrast, the temporary decline in scientific output between 2019 and 2021 may be related to the impacts of the COVID-19 pandemic and the weakening of environmental policies in Brazil. Contributing factors include the weakening of the Action Plan for the Prevention and Control of Deforestation in the Legal Amazon (PPCDAm), the suspension of the Amazon Fund in 2019 (Marcovith & Pinsky, 2020), and the weakening of environmental agencies such as the Brazilian Institute of Environment and Renewable Natural Resources (IBAMA) and the Chico Mendes Institute for Biodiversity Conservation (ICMBio) (MPI, 2023). Overall, although the temporal trend indicates a gradual increase in scientific production, especially over the past 5 years, only 41.48 % of the articles initially compiled directly addressed the role of Indigenous communities in Amazon biodiversity conservation.

### 3.2 Geographic distribution of studies

#### 3.2.1 Countries with the highest academic output

The analysis of the geographic distribution of studies by country showed that Brazil had the highest number of publications (n = 26), followed by Ecuador (n = 25) and Peru (n = 22). Bolivia accounted for 10 articles, Colombia for 7, French Guiana for 3, and Venezuela for 1 (Figure 3, Table S1). These differences may be associated with differences in the legal and institutional frameworks governing territorial rights and Indigenous participation in each country. Which directly influences research agendas, analytical focus, and the extent to which local communities are involved in scientific studies (Athayde et al., 2025; Ortiz et al., 2025).

**Figure 3.**
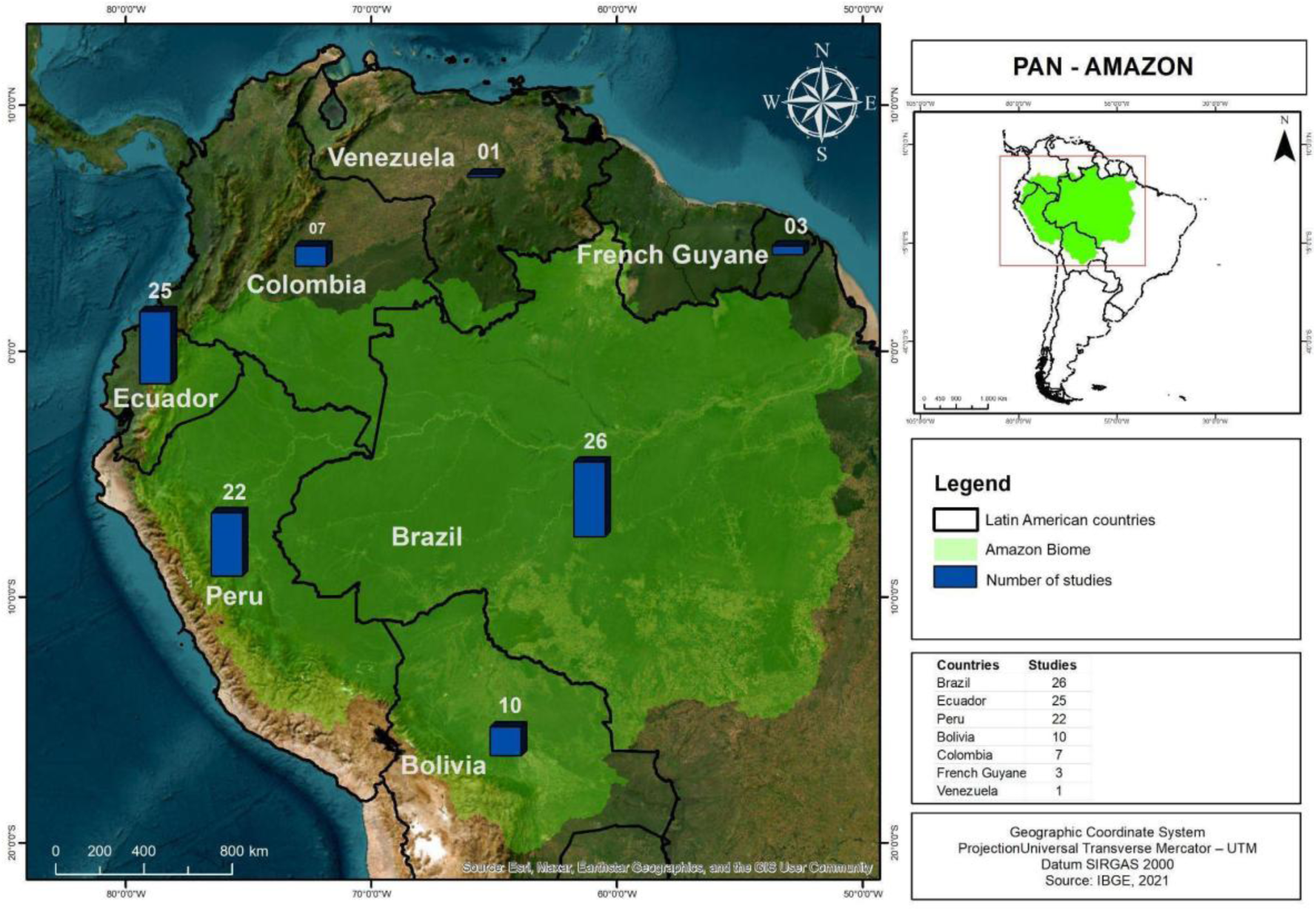
Geographic distribution map of studies addressing the participation of Indigenous peoples in Amazon biodiversity.

The Brazilian Amazon stood out as a major center of scientific production. Brazil contains approximately 60% of the Amazon biome, harbors exceptionally high biodiversity, and plays a key role in regional and global ecosystem and climate processes (Guayasamin et al., 2024; Levis et al., 2024; Guerrero-Moreno et al., 2025; Flores et al., 2024). This prominence may also reflect Brazil’s relatively robust legislative framework, particularly the National System of Protected Areas (SNUC), which organises protected areas and creates institutional conditions favourable to scientific research within legally recognised territories.

Collaborative conservation programs such as the Amazon Region Protected Areas Program (ARPA) have also played a major role in protecting millions of hectares of forest through international funding and community participation (Brazil, 2018). Another relevant framework is the National Policy for the Sustainable Development of Traditional Peoples and Communities (PNPCT), which recognizes and strengthens traditional ways of life and the sustainable use of natural resources, helping to integrate biodiversity conservation with sociocultural rights (Brazil, 2007).

Most studies identified in Brazil were conducted on legally recognised Indigenous Lands, including the Kayapó, Baniwa, and Zó’é territories. Studies were also carried out in protected areas such as Biological Reserves and Extractive Reserves, which, although not legally classified as Indigenous Lands, were included because they are inhabited by Indigenous communities or embedded within territories of traditional use (Toledo et al., 2015). Research in these areas has focused mainly on management and conservation strategies linked to participatory Indigenous practices, including environmental monitoring and participatory biodiversity inventories. One emblematic example is the leading role of Indigenous communities in monitoring flooded forests affected by the Belo Monte dam (Utsunomiya et al., 2024; Quaresma et al., 2025). Another is the ethnobotanical survey conducted among the Baniwa in the Rio Negro region near Manaus, which documented traditional knowledge of campinarana landscapes and the use of forest resources (Abraao et al., 2008).

Despite these advances, major challenges remain in the Brazilian Amazon, including rising illegal deforestation, mining, and land conflicts, which frequently threaten both biodiversity and the rights and agency of Indigenous peoples (Rorato et al., 2021; Mataveli et al., 2022). Even with a relatively well-established environmental framework, stronger public policies are still needed, particularly for land demarcation, because formal recognition of Indigenous territories supports both the maintenance of traditional knowledge systems and biodiversity conservation (Gonçalves et al., 2021).

In Ecuador, Indigenous ancestral knowledge receives particular recognition in the Constitution, which is widely regarded as pioneering for incorporating the rights of nature into the legal system (Pacheco, 2012). This framework recognizes nature, or *Pacha Mama*, as a subject of rights, a concept deeply rooted in Indigenous worldviews (Pacheco, 2012; Barcellos et al., 2023). Originating from Quechua, *Pacha Mama* can be translated as “Mother Earth” or “Mother Cosmos”, reflecting a view of the planet as living and interconnected (Pacheco, 2012). Its incorporation into the Constitution resulted from a long process of Indigenous struggle and advocacy. It marked a major advance in environmental law by recognizing nature as a rights-bearing subject rather than merely an object of exploitation (Barcellos et al., 2023). Ecuador has also implemented initiatives such as the *Socio Bosque* program, which provides economic incentives to Indigenous communities that maintain their forests, thereby strengthening partnerships between the state and traditional populations in sustainable natural resource management (Koning et al., 2010; Holland et al., 2014).

Among the most studied territories in Ecuador were Yasuní National Park and the Sumaco Biosphere Reserve, both in the Ecuadorian Amazon. Research focused on these areas may reflect the overlap between protected areas and Indigenous territories, thereby favouring collaborative conservation and sustainable management initiatives (Holland et al., 2014). In this context, interactions between public policies and community practices have produced positive outcomes for forest conservation (Bremner & Lu, 2006).

In the Peruvian Amazon, studies were conducted mainly in Manu National Park, the Maijuna-Kichwa Regional Conservation Area (MKRCA), and the Imiría Regional Conservation Area. This pattern may be linked to initiatives under Peru’s National System of State-Protected Natural Areas (SINANPE), which promotes the conservation of biological diversity and ecosystem services nationwide.

One relevant example is the collaboration between the Indigenous Shipibo community of Nuevo Saposoa and the government platform *Geobosques*, designed to combat illegal logging (World Resources Institute, 2017). Through this partnership, Shipibo community members received training to operate drones and conduct satellite-based monitoring, enabling them to capture detailed images of threatened forest areas. These data are shared with local authorities, strengthening enforcement and increasing Indigenous autonomy in forest conservation, natural resource protection, and territorial defense (González et al., 2023).

By contrast, Bolivia, Colombia, French Guiana, and Venezuela were less represented in the literature analyzed. Although these countries also contain important portions of the Amazon and high biological diversity, they accounted for relatively few studies. This lower representation may be associated with funding limitations, logistical challenges, and structural inequalities in the distribution of research resources and scientific infrastructure (Barlow et al., 2016; Fernandes et al., 2025).

#### 3.2.2 Comparative analysis between the most productive countries and the most studied *countries*

We observed a clear mismatch between where research is conducted and where academic leadership is based. Although Brazil, Ecuador, and Peru accounted for most of the studies, academic leadership was concentrated in the Global North, especially the United States (n = 17), the United Kingdom (n = 11), and Spain (n = 8) (Figure 4).

**Figure 4.**
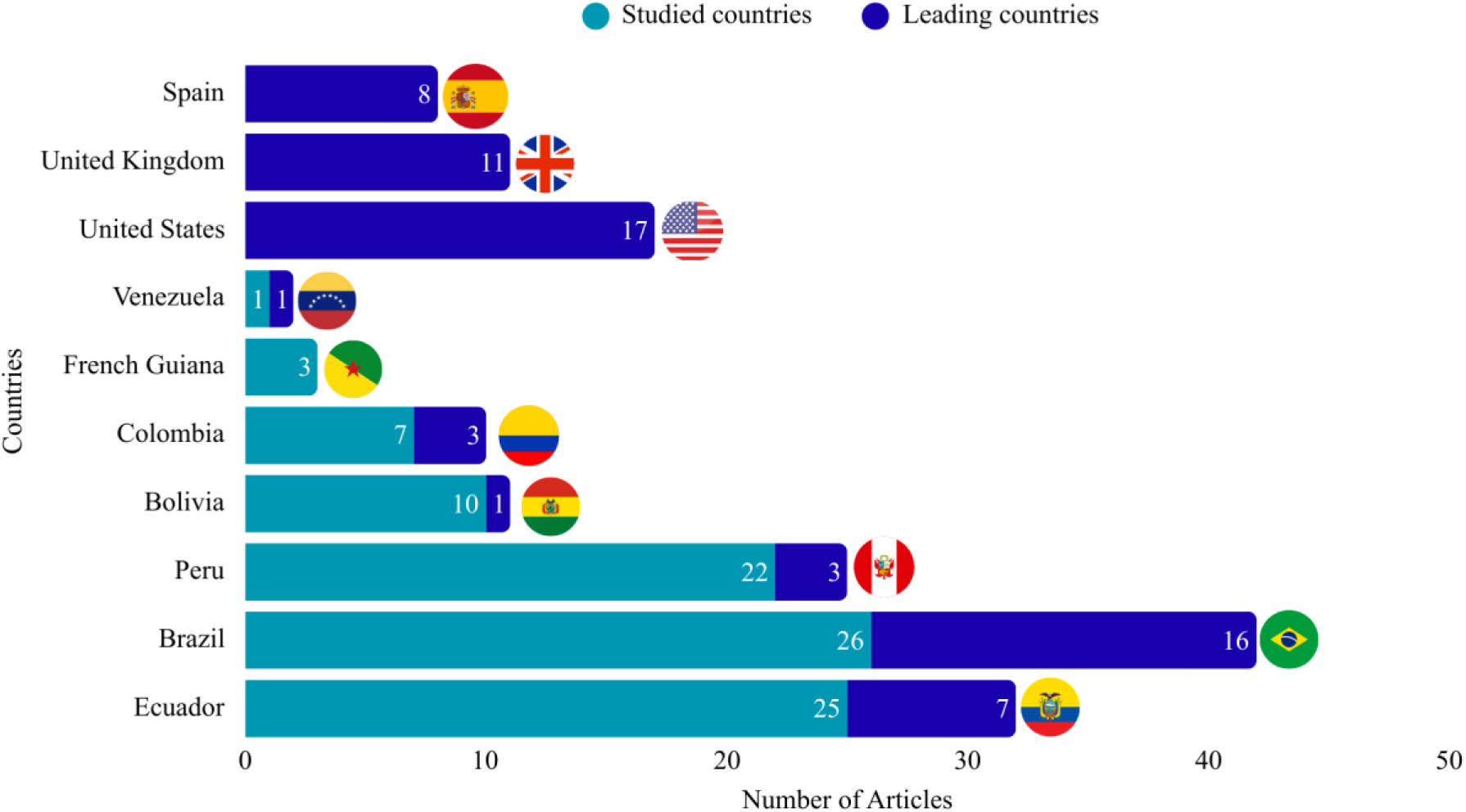
Most studied countries and countries leading academic production on Indigenous peoples’ participation in Amazon biodiversity conservation.

Taken together, these results show that although the Amazon is the main focus of study, much of the knowledge produced is driven by institutions outside the region. This disparity may reflect the greater availability of scientific infrastructure, funding, and international collaboration networks in the Global North, which facilitate both the development and publication of research on conservation and Indigenous peoples in the Amazon (Guerrero-Moreno & Oliveira-Junior, 2024; Da Silva et al., 2025).

### 3.3 Main institutional affiliations and Indigenous co-authorship in Amazon biodiversity *conservation*

The analysis of authors’ institutional affiliations showed that a relatively small group of institutions plays a central role in scientific production on biodiversity conservation involving Indigenous peoples in the Amazon (Figure 5). The most represented institutions were the National Institute of Amazonian Research (INPA) and Amazonas State University (UEA), with 13 studies each (13.82%). These were followed by the Universitat Autònoma de Barcelona (n = 12; 12.76%), the University of East Anglia (n = 10; 10.63%), George Mason University (n = 9; 9.57%), the University of Manchester (n = 7; 7.44%), Wageningen University (n = 7; 7.44%), the University of Leeds (n = 6; 6.38%), and the University of São Paulo (n = 6; 6.38%). The Instituto Socioambiental (ISA), a non-governmental organization with a strong socio-environmental presence in the Amazon, also stood out, appearing in 6 publications (6.38%) (Figure 5).

**Figure 5.**
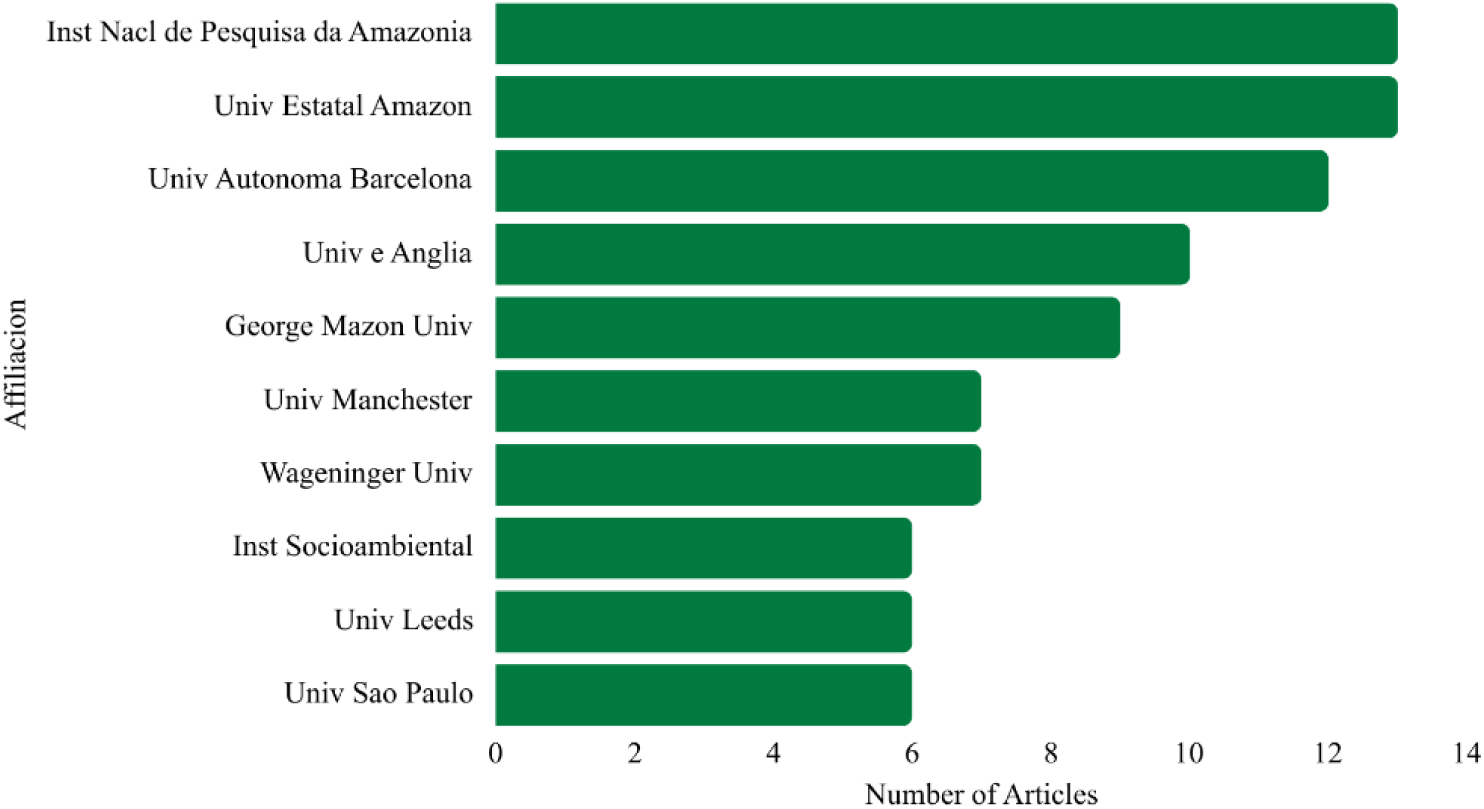
Institutions involved in producing studies on Indigenous peoples’ participation in Amazon biodiversity conservation.

The strong presence of Amazon-based institutions such as INPA and UEA reflects their longstanding role in advancing research on biodiversity and on the relationships between science and traditional knowledge in the region. INPA has become one of the leading institutions dedicated to the study of Amazonian biodiversity and the training of researchers in ecology and conservation (Manzi et al., 2016). Likewise, Amazonas State University has played an important role in the academic training of Indigenous students and in strengthening initiatives centered on intercultural education and the valuing of traditional knowledge, thereby bringing scientific production closer to the sociocultural realities of the region (Estácio & Almeida, 2016).

Among the initiatives linked to this process is a CAPES-approved project aimed at training teachers, preferably Indigenous teachers, to work in their own communities. This initiative promotes the integration of traditional and academic knowledge, strengthening intercultural education and expanding Indigenous participation in teaching and research (FAPEAM, 2025).

Despite these initiatives, authorship analysis showed that Indigenous participation in the scientific literature remains limited (Guerrero-Moreno et al., 2024). Only six of the studies identified included Indigenous co-authors. This highlights a discrepancy between the central role of Indigenous peoples in Amazon biodiversity conservation and their representation in formal academic production. Such underrepresentation may be linked to historical inequalities in access to formal education and scientific training, as well as to the limited implementation of public policies that support Indigenous academic training at advanced levels (Barbalho et al., 2025). The historical marginalization of traditional knowledge systems, often treated as peripheral to Western science, further contributes to the underrepresentation of Indigenous researchers in the scientific literature (Levis et al., 2024). Another source of these asymmetries is structural inequality in global scientific production. Although much of the research is conducted in the Amazon, many editorial networks, funding structures, and research centers remain concentrated outside the region, particularly in the Global North (Ordóñez-Matamoros et al., 2011).

Research investment also remains unevenly distributed within Brazil. Although the Brazilian Amazon is one of the most studied regions, financial resources for biodiversity research are not equitably distributed across regions. Investments linked to the Brazilian National Institutes of Science and Technology (INCT) have contributed significantly to advancing biodiversity knowledge, but resources remain concentrated elsewhere. For example, the Southeast has the largest number of funded research networks, whereas in the North, only Manaus and Belém host networks dedicated to Amazonian biodiversity research (Resende et al., 2025; Athayde et al., 2025). Although 13 INCT networks are currently active in the North, research specifically focused on Amazonian biodiversity remains concentrated in these two cities.

Efforts to reduce these inequalities include the Amazon Biodiversity Synthesis Network (INCT-Symbian), launched in Belém in December 2023. The network brings together researchers from multiple institutions and regions to consolidate Amazon biodiversity research and strengthen interdisciplinary collaboration, including knowledge co-production with Indigenous communities (Resende et al., 2025). Practical examples of this approach include *Human & Planet-Xingu Biodiversity Monitoring*, developed with Conservation International (CI), which involves Panará leaders and researchers from the Federal University of Pará (UFPA). This initiative strengthens community autonomy of Indigenous territories by integrating scientific monitoring with traditional knowledge. Another recent initiative is the Advanced Research-Action Centre for Amazonian Ecosystem Conservation and Recovery (CAPACREAM), launched in 2025 to promote research on Amazonian socio-biodiversity and the recovery of degraded ecosystems. In one of its associated projects, *SocioBIO,* it integrates traditional knowledge and academic science in conservation and ecological restoration strategies in the Amazon region (CAPACREAM, 2025). Despite these initiatives, further investment and stronger public policies for Indigenous academic training are crucial for supporting the diversity of perspectives in scientific knowledge production and biodiversity conservation in Indigenous territories (Flores et al., 2024).

### 3.4 Contributions of Indigenous communities to biodiversity conservation in the Amazon

The literature analyzed revealed four main categories of Indigenous community contributions to biodiversity conservation in the Amazon (Figure 6). Community-based monitoring and management was the most prominent category (n = 39; 41.48%), followed by cultural practices and ecological knowledge (n = 36; 38.29%) and management and sustainable use of natural resources (n = 18; 19.14%). In contrast, habitat conservation and restoration were the least documented.

**Figure 6.**
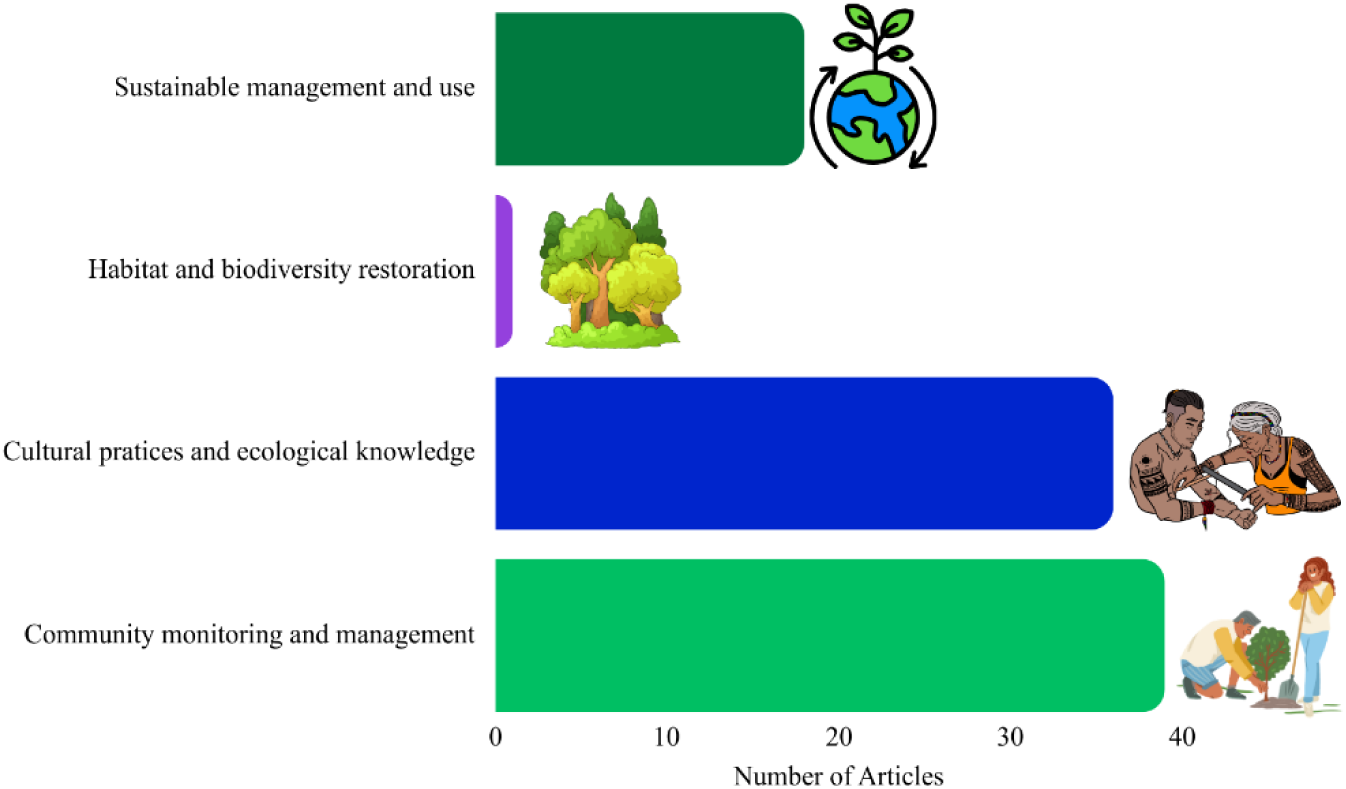
Main contributions of Indigenous peoples to biodiversity conservation in the Amazon.

These results highlight the leading role of Indigenous peoples in generating and interpreting environmental information. These initiatives include systematic tracking of species abundance, monitoring of environmental change, and territorial management grounded in local knowledge. Participatory monitoring can improve the reliability of environmental assessments and strengthen conservation decision-making (Walker et al., 2004; Silva et al., 2024). A relevant example is the work developed by Conservation International (CI) in partnership with Indigenous communities. In the Kayapó territory of A’Ukre, collaborative projects assessed and documented threatened species by combining scientific methods with traditional knowledge (Zimmerman et al., 2001). Likewise, initiatives promoted by the Instituto Socioambiental (ISA) through the *Y Ikatu Xingu* project have trained socio-environmental agents around the Xingu Indigenous Park, helping to strengthen territorial protection and restore springs and riparian forests threatened by agricultural expansion.

The category of cultural practices and ecological knowledge reflects how traditional communities perceive and interact with nature through knowledge systems grounded in cultural values, traditional practices, and accumulated experiences across generations. These systems guide the sustainable use of natural resources and play a fundamental role in maintaining biodiversity (Thevenin et al., 2019; Fernandes et al., 2021). Their importance is well documented in ethnobotanical studies showing close links among culture, biodiversity, and the use of plants for medicinal, food, and spiritual purposes (Prance, 1997). One example is Pilnik et al. (2023), who worked with the Huni Kuin people to document the cultural values associated with food plants and highlighted both the utilitarian and symbolic dimensions of plant species.

Studies conducted in partnership with Indigenous communities have also documented fauna through inventories and observations. These include records of Indigenous traditional knowledge of birds (Greeney et al., 2018), surveys of fish species (Bogotá et al., 2024), and studies of mammals (Stafford et al., 2016). Other works, including Levis et al. (2018), Thomas et al. (2010), Robert et al. (2012), and Abraao et al. (2008), further reinforce the role of Indigenous traditional knowledge in the use and sustainable management of landscapes and ecosystems.

The category management and sustainable use of natural resources include practices that promote balanced resource use and maintain productive systems compatible with environmental conservation. Strategies documented in the literature include traditional agroforestry systems, such as *chakra* systems in the Ecuadorian Amazon (Rodriguez et al., 2021), as well as sustainable fishing and hunting practices regulated by community norms that define permitted periods, locations, and species (Ohl-Schacherer et al., 2007; Constantino et al., 2008). These practices have generated positive outcomes for both biodiversity conservation and food security. Furthermore, the Indigenous Agroforestry Agents program, developed by the Comissão Pró-Índio do Acre (CPI-AC), provides long-term training to strengthen territorial management and the sustainable management of fauna and flora in Indigenous territories. The program was created in response to requests from Indigenous leaders for support in managing the use of wild fauna in Kaxinawá territory.

Environmental conservation and restoration were the least-represented category. It includes initiatives focused on ecosystem restoration, the creation and management of protected areas, and the recovery of degraded habitats. The only study identified in this category described the return of the giant otter (*Pteronura brasiliensis*) to the Içana River in Baniwa territory, highlighting the potential of Indigenous territorial conservation strategies to support the recovery of threatened species (Pimenta et al., 2018).

Overall, these results indicate that despite the ecological relevance of Indigenous territories and the active role of Indigenous communities in biodiversity conservation, traditional knowledge remains underrepresented in parts of the scientific literature. The absence of these perspectives can limit understanding of Amazonian socioecological systems and constrain the development of more inclusive and effective conservation strategies. In this sense, integrating traditional knowledge and academic science has been recognized as a key approach for developing fairer and more effective environmental policies, strengthening Indigenous autonomy, and improving the ecological information used in environmental governance (Levis et al., 2024).

### 3.5 Taxa assessed and ecosystem types studied in biodiversity conservation in Indigenous territories in the Amazon

Research conducted in Indigenous territories across the Amazon covers a wide diversity of taxa, reflecting the biological richness of these ecosystems. The most frequently investigated groups were plants, mammals, fish, birds, and insects, chosen because of their ecological, cultural, and economic importance to Indigenous communities (Figure 7).

**Figure 7.**
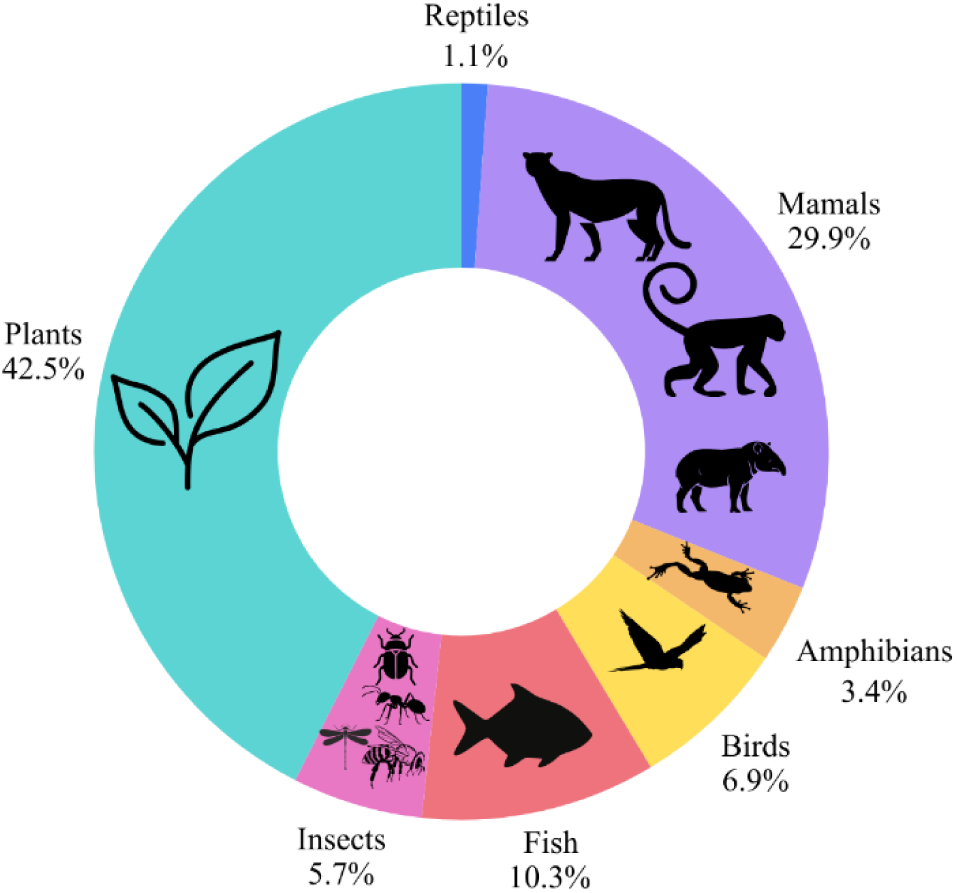
Taxa or taxonomic groups assessed in studies involving Indigenous peoples and biodiversity conservation in the Amazon.

Among the taxa investigated, plants were the most frequently studied group (n = 37; 42.5%). Most studies focused on cultivated species (Arango-Ulloa et al., 2009; Thomas et al., 2010), medicinal and food plants (Pilnik et al., 2023; Guachamin-Rosero et al., 2022), and species associated with traditional agroforestry systems (Oldekop et al., 2013; Torres et al., 2024; Robert et al., 2012). For Indigenous communities, plants are central to subsistence, traditional medicine, and cultural and religious practices (Leret-Cámara et al., 2019). These studies were conducted in territories including the Sumaco Biosphere Reserve (Ecuador), the Kaxinawá and Baniwa Indigenous Lands (Brazil), and Kichwa communities in Ecuador. These data combined workshops, interviews, and floristic inventories.

Mammals were the second-most-studied group (n = 26; 29.9%). Many studies were conducted in the Kaxinawá and Kayapó Indigenous Lands (Brazil), the Cofán territory (Ecuador), the Tsimane territory (Bolivia), and Manu National Park (Peru). Their prominence reflects their importance to Indigenous subsistence, both as primary hunting resources and as biological indicators in wildlife monitoring (Iwamura et al., 2014; Wallace et al., 2012). The main methods included camera trapping, semi-structured interviews, and assessments of hunting sustainability.

Fish ranked third (n = 9; 10.3%). Studies were conducted in areas such as Yasuní National Park (Ecuador) and Maijuna territory (Peru), using inventories and population monitoring (Bogotá et al., 2024), as well as analyses of the relationships between fish and community use of aquatic resources (Mora et al., 2018; Oviedo et al., 2017). These studies combined capture-based methods, interviews, and modelling approaches.

Birds were relatively underrepresented (n = 6; 6.9%). Research was conducted in the Kaxinawá Indigenous Lands (Brazil), Amacayacu National Park (Colombia), and the Yasuní Biosphere Reserve (Ecuador), with a primary focus on population monitoring, management, and conservation practices. The most common methods were visual and acoustic surveys, combined with interviews on local perceptions and practices associated with birdlife (Arcilla et al., 2025; Greeney et al., 2018).

Insects were among the least represented groups (n = 5; 5.7%). These studies were conducted mainly in Kichwa communities and the Yasuní Biosphere Reserve in Ecuador, as well as Extractive Reserve areas in Brazil. They focused especially on edible insects such as beetles and ants, which play important roles in the diets and food cultures of many Amazonian communities (Gasca-Álvarez et al., 2022). The studies combined traditional knowledge, specialized collection techniques, and ethno-entomological analyses. Some also pointed to the potential of insects as key organisms for ecotourism and citizen science, contributing to the appreciation of local biodiversity (Guerrero-Moreno et al., 2024; Ribeiro et al., 2025).

Among vertebrates, amphibians (n = 3; 3.4%) and reptiles (n = 1; 1.1%) were the least studied groups. Research on both was conducted on the Kayapó Indigenous Land (Brazil) and in Manu National Park (Peru), primarily using these organisms as indicators of environmental impacts (Oldekop et al., 2013).

Overall, studies conducted in Indigenous territories across the Amazon show a strong predominance of organisms associated with terrestrial ecosystems, to the detriment of aquatic organisms. This reflects a widely documented trend in biodiversity research, which tends to focus on plants and vertebrates while leaving other taxonomic groups underrepresented (Troudet et al., 2017; Mammola et al., 2020). Aquatic ecosystems such as streams, rivers, and lakes remain underexplored despite their ecological and cultural importance for Amazonian populations (Castello et al., 2018; Lopes et al., 2021). This gap is also shaped by the logistical complexity of research in Amazonian aquatic environments, which often requires specialized equipment, longer expeditions, and extensive travel to remote areas (Carvalho et al., 2023; Arantes et al., 2019).

In addition to these structural barriers, the limited inclusion of Indigenous communities in research processes may further constrain advances in knowledge about these ecosystems. The historical marginalization of traditional knowledge within Western science has contributed to the underuse of local knowledge that could substantially expand understanding of Amazonian biodiversity (Levis et al., 2024; Athayde et al., 2025). Given the central role of aquatic ecosystems in the subsistence and territorial organization of Indigenous communities across the Amazon, expanding research in these environments is essential for a more integrated understanding of biodiversity dynamics in the world’s largest river basin.

### 3.6 Indigenous perspectives on scientific production and pathways for change

The results of this study reveal low Indigenous representation in scientific authorship and a concentration of academic leadership in institutions based in the Global North. Because these asymmetries cannot be understood solely from an external academic perspective, we shared this manuscript with Panará Indigenous researchers and representatives of other Amazonian peoples, inviting them to contribute to the critical interpretation of the findings. These contributions broaden the debate by incorporating Indigenous researchers’ experiences and perceptions of how science has been produced in their territories.

The Indigenous researchers’ narratives help illuminate how these inequalities are expressed in practice and what changes are needed to support genuinely collaborative science. In this sense, this section represents an exercise in knowledge co-production, in which different knowledge systems engage in dialogue to interpret the existing scientific literature critically. These contributions were incorporated as direct narratives, while formally recognizing all Indigenous researchers as co-authors of the manuscript.

For the Panará representatives, the small number of scientific studies that include Indigenous people as authors reflects a long history of inequality in knowledge production. In many cases, Indigenous peoples participate in fieldwork and contribute deep knowledge of territories, species, and forest dynamics. Yet, these contributions do not always appear formally in the authorship of scientific publications (Haelewaters et al., 2021). Participants also interpreted the predominance of institutions from the Global North as resulting from structural inequalities in science, particularly in access to funding, infrastructure, and international research networks. At the same time, this context reinforces the importance of strengthening research institutions in the Amazon and Brazil, and of expanding the leadership of local researchers, especially Indigenous researchers.

Based on their experience in the biodiversity monitoring project in the Panará territory, participants identified biodiversity monitoring, water quality, environmental change, dynamics of fauna and flora, and the documentation of traditional knowledge related to plants and forest relationships as priority research themes in Indigenous territories. Participants further emphasized that science in Indigenous territories must advance in dialogue, transparency, and participation. Communities should be involved from the planning stage, participate in field activities, and have access to results in practical and accessible formats. Initiatives that promote the training and direct participation of Indigenous researchers are likewise essential for strengthening autonomy in the production of knowledge about their own territories.

For the Indigenous researcher Tyson Ferreira-Sateré: “*Although we are beginning to occupy some spaces within scientific culture through affirmative action policies and institutional invitations, this participation is still limited and often not accompanied by recognition or trust from other scientists and institutions. We are frequently seen as incapable or as holders of incomplete knowledge, reduced to myths, beliefs, or folklore. Yet that same knowledge, when appropriated by external researchers, becomes validated as universal science, rendering Indigenous leadership in scientific production invisible. As an Indigenous person, I interpret this predominance as a form of scientific colonialism, in which Indigenous peoples are treated as objects of study rather than as subjects of knowledge. Researchers collect data and traditional knowledge and publish them without ensuring that this knowledge returns to the communities or is properly recognized at its source. In light of this, a profound paradigm shift is needed: moving from an extractive logic to a collaborative one. Science should not be conducted ‘about’ Indigenous peoples, but with them. Research questions should emerge from the needs of the territories, and results should be returned to communities in accessible formats*” (Ferreira-Sateré, personal communication, 2026).

For the Indigenous researcher Amanda Kumaruara: “*When I see that few scientific studies include Indigenous people as authors, I understand this as a reflection of a long process of invisibilities. We have always been present in research, in the field, in the territory, sharing our knowledge, but we are rarely recognized as producers of that knowledge. This reveals a structural inequality in the way science is built*” (Kumaruara, personal communication, 2026). She further emphasized that the predominance of research led by countries from the Global North reinforces an unequal relationship in which Indigenous territories are studied, and Indigenous knowledge is used. At the same time, recognition remains concentrated outside these contexts. “*Too often, people speak about us without us. I believe science needs to change the way it relates to Indigenous peoples, moving away from a logic of knowledge extraction and towards joint construction. This means including Indigenous people from the very beginning of research, ensuring genuine co-authorship, and respecting different ways of producing knowledge*” (Kumaruara, personal communication, 2026).

For the Indigenous researcher Yuri Kuikuro: “*As an Indigenous researcher, I observe that scientific production still needs to recognize more broadly the knowledge built by Indigenous peoples. Our knowledge comes from a continuous relationship with territory, from observing nature, and from the transmission of knowledge across generations, integrating cultural, spiritual, and environmental dimensions. I also see that there are still too few Indigenous people in decision-making roles within academic and scientific spaces. It is essential to expand Indigenous participation in these contexts because it makes no sense to speak about us without our presence. The inclusion of Indigenous researchers strengthens science, making it more ethical, more diverse, and more connected to the realities of the territories*” (Kuikuro, personal communication, 2026). He also stressed that Indigenous peoples are not homogeneous but differ in culture, language, and social organisation. For this reason, it is necessary to expand the presence of different Indigenous peoples in academia, ensuring that multiple perspectives are represented in knowledge production. “*In the Amazon and in other Indigenous territories, there is a great diversity of peoples who observe and understand nature through their own experiences and through the natural cycles lived in their territories. These forms of knowledge contribute to expanding scientific understanding of the environment, biodiversity, and the relationships between society and nature. I believe that strengthening Indigenous presence in universities and research is essential for building a more inclusive science, one that respects cultural differences and values different ways of observing and understanding the world. In this way, scientific production can become a space for dialogue among knowledge systems, contributing to more sustainable and socially just pathways*” (Kuikuro, personal communication, 2026).

Taken together, these narratives show that the patterns identified in the bibliometric analysis reflect concrete experiences lived by Indigenous peoples in the Amazon in relation to scientific production. Their contributions reinforce the need for structural transformation in science, grounded in inclusion, recognition, and knowledge co-production, and for aligning scientific practices more closely with the sociocultural realities of Indigenous territories.

## 4. Conclusion

This study highlights both the potential and current limitations of Indigenous participation in scientific production on Amazon biodiversity conservation. Over nearly three decades, research on this topic has gradually expanded, underscoring the importance of co-producing knowledge with Indigenous communities. Still, Indigenous representation remains limited, despite these communities holding complex knowledge accumulated across generations and acting as key observers, interpreters, and managers of their environments.

Strengthening the co-production of traditional and scientific knowledge is essential for advancing research across terrestrial and aquatic ecosystems. At the same time, the contrast between the most-studied countries and those leading academic production reveals the predominance of researchers from the Global North, who hold most institutional affiliations and positions of scientific leadership, even when research is conducted in Amazonian territories. This pattern reflects limited recognition of Indigenous leadership, the underrepresentation of researchers from the Global South, and structural and logistical barriers that continue to constrain local scientific autonomy. A more equitable science, aligned with the needs of Indigenous territories, can strengthen integrated knowledge systems, recognize Indigenous roles in biodiversity protection, and help address historical gaps.

Collaborative research in Indigenous and traditional territories also raises growing concerns about data governance, including where research data go and who controls their access, responsibility, and use. Beyond compliance with legislation on traditional knowledge associated with genetic heritage, Indigenous leaders in some countries have emphasized the relevance of the CARE Principles for Indigenous Data Governance, centered on collective benefit, authority to control, responsibility, and ethics. These principles provide an important starting point, but must be further discussed.

Overall, our findings reinforce the need for more symmetrical and collaborative approaches to biodiversity conservation research. More inclusive and equitable conservation science will depend not only on recognizing Indigenous knowledge, but also on supporting Indigenous leadership, shared authority, and fairer collaboration in the production and governance of knowledge.

## Statements

### Author contributions

Jady Vivia Almeida da Silva Santos, Francieli F. Bomfim, Thaisa Sala Michelan, Karina Dias-Silva, Luciano Fogaça de Assis Montag and Leandro Juen conceived the ideas and designed methodology. Jady Vivia Almeida da Silva Santos, Yan Campioni, Everton Cruz da Silva, Kiarasy Kaiabi Panara, Korakoko Panara, Sewa Panara, Sakre Panara, Karapow Panara, Kwakore Panara, Sopoa Panara, Nhasykiati Panara Pâssua Pri Panara, Pente Panara, Tepakriti Panara, Tyson Ferreira-Sateré, Amanda Kumaruara, Yuri Kuikuro, Antonio Ramyllys Oliveira Costa, Lais Sarlo, Bruno Coutinho, Renata Pinheiro and Rafael de Araújo collected the data. All authors analysed the data. Jady Vivia Almeida da Silva Santos, Francieli F. Bomfim, Thaisa Sala Michelan, Karina Dias-Silva, Luciano Fogaça de Assis Montag and Leandro Juen led the writing of the manuscript. All authors contributed critically to the drafts and gave final approval for publication.

### Data availability statement

Data will be available upon reasonable request.

## Acknowledgements

We are grateful to Conservation International (CI) for financial support through the authors’ grant JVASS (Donation No. CI-115395). This study was also conducted within the framework of the Programa de Pesquisa em Biodiversidade da Amazônia Oriental (PPBio AmOr - process 441257/2023–2) and was supported by the Coordenação de Aperfeiçoamento de Pessoal de Nível Superior – Brazil (CAPES), Financing Code 001. The authors are grateful to the National Council for Scientific and Technological Development (CNPq) for scholarships and grants to PERP (process 173670/2023-7), GBV (process 382645/2025-1), EGP (process 380068/2026-5), and Research Productivity Grants to LFAM, RAMS, LJ, TSM, MPDS and DJR (processes 302881/2022-1, 304710/2019-9, 311835/2023-6, 305551/2023-1, and 312407/2022-0). We appreciate the support of the projects INCT Sínteses da Biodiversidade Amazônica (process 406767/2022–0), Advanced Research-Action Center for the Conservation and Recovery of the Amazon Ecosystem (CAPACREAM - process 444350/2024-1), and Programa de Pesquisa de Longa Duração (PELD-AmOr - process 445970/2024–3). We thank the field team from the Federal University of Pará, as well as the CI team, for their logistical and technical support during the expedition days, and the Panará Indigenous community, who welcomed us with great affection to their territory.

---

**Table S1.** Amazonian countries and their respective Indigenous territories, together with the number of articles per country included in this study.

| Amazonian Countries | Indigenous Territories with recorded Studies | Number of articles |
| --- | --- | --- |
| Brazil | TI Kayapó; TI Zoé; TI Gavião, TI Kaxinawá; TI Baniwa; TI Santeré - Mawé; Reserva Biológica do Gurupi; Reserva Extrativista Tapajós-Arapiuns | 26 |
| Ecuador | Parque Nacional Sumaco; Cofán, Kichwa; Shuar; Achuar: Estação Biológica Timburí Cocha (TCBS); Parque Nacional Yasuní; Pakayaku | 25 |
| Peru | Parque Nacional de Manu; Conservação Regional Maijuna-Kichwa (MKRCA); Área de Conservação Regional Imiría; | 22 |
| Bolivia | Tsiname; Parque Nacional Isiboro-Sécure; Reserva da Biosfera Pilon-Lajas | 10 |
| Colombia | Reserva Indígena Uitoto de Monochoa; Arhuaco; Cubeo; Parque Nacional Amacayacu | 7 |
| French Guiana | Wayãpi; Makushi e Wapishana | 3 |
| Venezuela | Sierra Maigualida | 1 |

